# Uncovering cross-signal interactions in single-cell and spatial omics

**DOI:** 10.64898/2026.11.03.763866

**Authors:** Jeremie Sasson, Ben Niasoff, Jonathan Karin, Yuval Ezraty, Mor Nitzan

## Abstract

Individual cells simultaneously encode multiple biological signals in their gene expression profiles, including spatial position, circadian phase, and disease state. These signals can be entangled in complex ways, including circadian patterns that vary along disease progression and cell cycle dynamics that shift along developmental trajectories. While methods to disentangle such signals have been recently developed, characterizing the interaction itself between multicellular signals along with its effects on tissue function, localizing cellular subpopulations which exhibit signal interactions, and identifying potential candidate genes which drive such interactions, all remain major challenges. We present CRISP which characterizes signal interactions in single-cell gene expression data based on the correspondence between the multiple, probabilistic signal-specific projections of the data. Across simulations with tunable couplings and four single-cell and spatial transcriptomic datasets, CRISP recovers diverse biological signal interactions and localizes them to informative cellular subpopulations and groups of associated genes; CRISP distinguishes healthy from fibrotic liver samples and localizes the loss of zonation–position coupling to individual fibrotic hepatocytes, resolves the asymmetric coupling of TGF-*β* response, EMT, and cell-cycle phase to specific cell subpopulations, and flags transcriptionally delayed hepatocytes in Plasmodium liver infection. Beyond quantifying and localizing signal interactions, CRISP can refine existing labels, uncover continuous cellular transitions and reveal condition-associated genes.

## 1 Introduction

The gene expression profiles of cells simultaneously encode multiple facets of their identity, ranging from cell type and tissue-of-origin to spatial location, temporal phase along developmental or circadian trajectories, and disease state [31, 30, 29, 28]. Single-cell RNA sequencing (scRNA-seq) has emerged as a powerful tool to dissect this multifaceted heterogeneity at the resolution of individual cells.

These distinct biological facets, or signals, are not merely co-expressed, but can be entangled within the same cell [31, 4, 18]. In the mammalian liver, for example, a subset of hepatocyte genes is both zonated and rhythmic, jointly encoding spatial location along the portal-central axis and circadian time [8]. Such joint encoding is not unique to the liver; cells generally co-encode signals related to their intrinsic state and extrinsic environment, including cell cycle phase, spatial context, lineage, and response to perturbations [8, 25, 24, 15, 7, 34, 9]. TGF-*β*, for example, drives cells through the epithelial-to-mesenchymal transition (EMT) [21, 23] while arresting the cell cycle [20]. Interactions between biological signals can directly influence tissue function. In the liver, zonal differences in drug-metabolizing enzyme expression are thought to underlie the zone-specific damage caused by many hepatotoxins [17], while circadian regulation modulates hepatotoxicity, as shown for acetaminophen in mice [14].

Recent studies have identified genes that encode two processes simultaneously by tailoring statistical models to given datasets and signal pairs; Droin et al. [8] fit mixed-effect models to mouse hepatocyte scRNA-seq, with separate spatial and temporal terms and a multiplicative interaction term, to classify each gene as zonated, rhythmic, both (additively or multiplicatively), or flat. In the same organ, Ben-Moshe et al. [3] developed a custom spatiotemporal analysis of zonal liver regeneration to isolate genes whose expression jointly tracks lobule position and time following acetaminophen injury. Such approaches yield deep biological insights, yet rely on design choices tightly coupled to a specific system. To handle the diversity of signals embedded in scRNA-seq data, several computational frameworks have been developed to disentangle signals of interest; SiFT [25] performs probabilistic filtering by projecting cells onto a similarity kernel that captures a target signal and subtracting the resulting projection, thereby exposing hidden cellular attributes. scPrisma [15] uses spectral template matching, leveraging topological priors over the structure of a signal (e.g., cyclic or linear) to enhance, filter, or reconstruct it. Several methods adopt deep generative or latent-factor approaches to disentangle sources of variation; Biolord [24] learns a disentangled latent representation of known and unknown attributes, TarDis [13] uses covariate-specific loss components in a self-supervised model to disentangle multiple categorical and continuous covariates into structured latent subspaces, and CellUntangler [7] embeds cells into multiple latent subspaces, each with a geometry tailored to a distinct signal. Latent-factor models such as f-scLVM [4] similarly decompose single-cell heterogeneity into annotated and unannotated factors, separating biological sources of variation from confounding effects.

While substantial progress has been made in signal disentanglement from single-cell data, characterizing how biological signals interact, and further which genes and cells encode such interactions, has not been generally addressed. The characteristics of such biological signal interactions can in turn effect collective tissue function and potential dysfunction upon disturbance.

Some forms of signal interactions can be detected solely based on the joint distribution of signal labels across cells. For example, if certain tissue locations contain cells at a single phase of the cell cycle while others contain cells across all phases, as is the case in the intestinal crypt-villus axis [22], then the two signals (spatial tissue locations and cell cycle progression) are coupled. Yet, even when the two distributions of signal labels across cells are independent, so that in our example the location of a cell is not indicative of its phase, the signals can still interact through changes in gene expression, as a gene expressed only at a specific location and a specific phase of day can tie the two signals together [8].

To address this gap, we introduce CRISP, Cross-Signal Representation Interaction via Similarity Projection, a framework that characterizes the interactions between a pair of biological signals through the correlation structure of the expression patterns they induce in single-cell data. For each signal, CRISP first isolates the expression profiles associated with it, by projecting the expression matrix onto a cell-cell similarity kernel that links cells with similar values of the labels associated with that signal. The interaction between the two signals is computed as the correlation between the two projected representations, where high correlation values reflect that the signals shape gene expression in overlapping ways. CRISP quantifies such signal interactions at three levels: a global tissue score for cross-dataset comparison, a cell-level score that localizes the signal interaction to individual cells, and a gene-level score that identifies genes that jointly encode both signals.

We validated CRISP on simulated data in which the level of interaction between two cyclic processes can be tuned in a controlled manner, and then applied it to three settings: the coupling between zonation and lobule position in healthy and fibrotic liver, between TGF-*β* dose and epithelial–mesenchymal transition, and between infection time and parasite developmental pseudotime in Plasmodium-infected hepatocytes. Beyond quantifying interactions between biological signals and identifying cells and genes that are associated with such interactions, CRISP can be used to refine existing labels, resolve continuous transitions that discrete categories mask, and reveal condition-associated genes.

## 2 Results

### 2.1 Quantifying signal interactions in single-cell data with CRISP

CRISP is a projection-based computational framework that quantifies the level of interaction between two different biological signals encoded in single-cell RNA-sequencing datasets and identifies the contribution of individual cells and genes to this interaction (Figure 1). CRISP takes as input a gene expression matrix *X* ∈ ℝ^*N* ×*G*^, with *N* cells and *G* genes, and two signals of interest; Each signal is represented by its per-cell label values 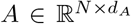 and 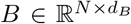, where *d*_*A*_ and *d*_*B*_ are the dimensions of the respective label spaces; for example, each cell’s spatial (*d* = 2) position and circadian phase (*d* = 1). Labels may be continuous or discrete, and the two signals need not share a type or topology (Methods 4.3).

**Figure 1:**
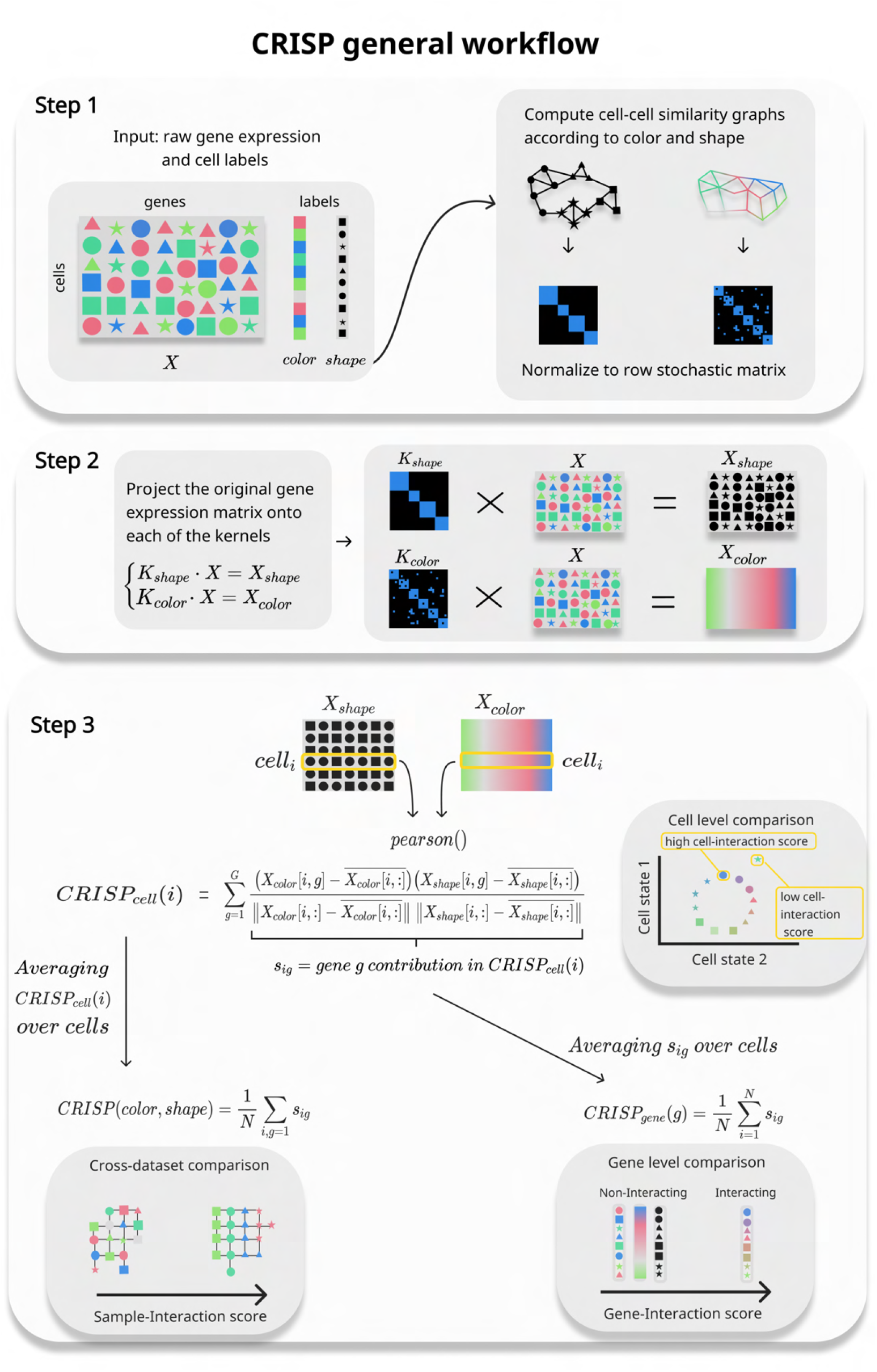
Overview of the CRISP framework. **Step 1** Input: a raw gene expression matrix *X* (*N* cells, *G* genes) and two cell-level labels (here, shape and color) that are used to define cell-cell similarity kernels. **Step 2** Projecting *X* onto each kernel gives *X*_shape_ and *X*_color_, retaining only the expression that co-varies with each signal. **Step 3** Row-wise Pearson correlation between *X*_shape_ and *X*_color_. Output: cell-interaction score CRISP_cell_(*i*), sample-interaction score CRISP(*color, shape*) and gene-interaction score CRISP_gene_(*g*).

We model two signals as interacting when they shape multicellular gene expression in similar ways, so that the organization of cells according to one signal (e.g. ordering of cells according to the cell cycle phases) is similar to the organization of cells according to the other signal (e.g. ordering of cells based on a spatial zonation pattern) [25, 24, 16]. Because a cell’s measured expression is a mixture of the contributions of multiple signals, CRISP first isolates the expression that varies with each signal by projecting the expression matrix onto a representation of the signal based on signal-specific cell-cell similarity kernel *K* ∈ ℝ^*N* ×*N*^ , whose entry *K*_*ij*_ reflects the similarity between cells *i* and *j* under that signal; for example, for the signal encoding the circadian phase, the kernel assigns high similarity to pairs of cells sampled at the same or adjacent times of day. Applying this separately to both signals yields two projections, *X*_*A*_ = *K*_*A*_*X* and *X*_*B*_ = *K*_*B*_*X*, both in ℝ^*N* ×*G*^. Each cell’s resulting projected expression is the average expression of its signal-specific neighboring cells. Such signal-specific expression smoothing preserves the signal-associated component in the gene expression, while remaining signal-independent variation is averaged out. For example, averaging the gene expression profile of a cell with cells sampled temporally nearby retains the expression pattern that varies with the time of day, while variation unrelated to it averages out.

Each cell *i* now has two associated expression profiles *X*_*A*_[*i*, :] and *X*_*B*_[*i*, :], each a vector of length *G* whose expression is smoothed across each of the two signals, respectively. For each cell, a cell-specific signal correlation score is computed as the correlation value between its two projected expression profiles, *X*_*A*_[*i*, :] and *X*_*B*_[*i*, :]. The cell-specific signal correlation score reflects the extent to which the two signals share an encoding, or are coupled, within that cell. Beyond the single cell, this approach aids in characterizing signal interaction at two further scales: averaged across all cells, characterizing how strongly the two signals interact dataset-wide; and split across genes, identifying the genes which carry the shared encoding.

Formally, CRISP builds a row-stochastic cell–cell similarity kernel *K* ∈ ℝ^*N* ×*N*^ for each signal, with *K*_*ij*_ the similarity between cells *i* and *j* under that signal (Methods 4.3). Since *K* is row-stochastic, each row of *KX* is a similarity-weighted average over a cell’s neighbors [25]. With *K*_*A*_ and *K*_*B*_ the kernels for signals *A* and *B*, respectively, the two projections are:

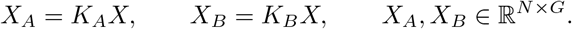

CRISP summarizes the congruence between *X*_*A*_ and *X*_*B*_ at three resolutions (Figure 1e–g). We write *X*_*A*_[*i*, :] for row *i* of *X*_*A*_, cell *i*’s projected expression onto signal *A*, and *X*[*i, g*] for the expression of gene *g* in cell *i*.

The **cell-interaction score** CRISP_cell_(*i*) for cell *i* is the Pearson correlation *r* between the two projections of cell *i*, taken across its *G* genes:

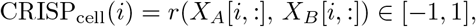

quantifying the contribution of cell *i* to the interaction between signals *A* and *B*. The **sample-interaction score** CRISP(*A, B*) between signals *A* and *B* is the mean of the cell scores over all cells in the dataset,

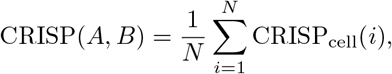

yielding a single, cross-dataset comparable measure of interaction.

The **gene-interaction score** splits each cell’s score among its genes. A cell-interaction score CRISP_cell_(*i*) is a Pearson correlation between each signal-specific gene expression, which is a sum over genes, 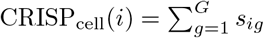, where the term for gene *g, s*_*ig*_, is:

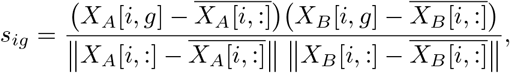

where 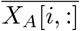 is the mean of row *i* over genes. The gene score CRISP_gene_(*g*) is gene’s *g* average contribution to the interaction, obtained by averaging the above term over cells:

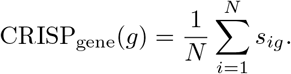

### 2.2 CRISP characterizes signal interactions at the sample, cell, and gene levels in simulated data

We validated CRISP on simulated data consisting of two interacting cyclic processes, *A* and *B*, with tunable coupling, alongside a third process *C* serving as a structured background. Specifically, every cell is assigned three cyclic labels, *θ*_*a*_, *θ*_*b*_, *θ*_*c*_ ∈ [0, 2*π*], defining a cyclic ordering of the cells according to processes *A, B*, and *C*, respectively. The phases of processes *A* and *B* are related by a fixed rotation, *θ*_*b*_ = *θ*_*a*_ + *φ*, while *θ*_*c*_ is drawn independently at random from a uniform distribution, providing a structured source of variation unrelated to the *A*-*B* interaction. Gene expression is then generated based on these labels: the simulated gene expression matrix *X* ∈ R^*N* ×*G*^, with *N* cells and *G* genes, comprises three gene programs, each a disjoint set of genes driven by one of the labels *θ*_*a*_, *θ*_*b*_, and *θ*_*c*_ (Methods 4.6.1). The expression profile of cell *i, x*_*i*_, is composed of the three programs,

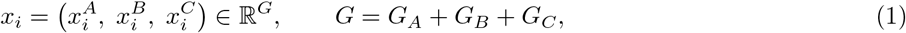

Where 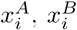, and 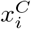 denote the restriction of *x*_*i*_ to the genes associated with each of the three programs *A, B*, and *C*, and *G*_*A*_, *G*_*B*_, *G*_*C*_ are the number of genes in each program. Because *θ*_*b*_ is a deterministic rotation of *θ*_*a*_, processes *A* and *B* are fully coupled in this initial setting: gene program *A* encodes *θ*_*b*_ as faithfully as *θ*_*a*_, and vice versa.

We control the interaction between the two signals using two independent parameters: (1) the cell permutation fraction *ρ* ∈ [0, 1], which jointly permutes the *B*-gene profiles and their *θ*_*b*_ labels in a fraction *ρ* of cells, interpolating between full *A*-*B* interaction (*ρ* = 0) and no interaction (*ρ* = 1); and (2) the number of interacting genes *G*_*AB*_, which encode both processes in every cell and form a fourth block of the expression profile (Methods 4.6.1, Figure 2a,b). We first evaluated the CRISP sample-interaction score. with the cell permutation fraction held at 1 - so that the *A* and *B* labels are independent - the CRISP sample-interaction score increases as the number of interacting genes *G*_*AB*_ increases, capturing shared transcriptional encoding beyond the information contained in the labels alone (Figure 2c). In contrast, a circular-regression baseline that fits the offset *φ* between the ground-truth labels *θ*_*a*_ and *θ*_*b*_ and reporting the mean cosine of the residuals (Methods 4.6.2) [19] remains constant and is structurally blind to the increasing number of interacting genes. This is due to the circular-regression score being computed based on the cell labels alone, which are unaffected by the number of interacting genes. As expected, increasing the permutation fraction from 0 to 1 when no interacting genes are present progressively weakens the *A*-*B* coupling, and both the baseline and the sample-interaction score decrease accordingly (Figure 2d).

**Figure 2:**
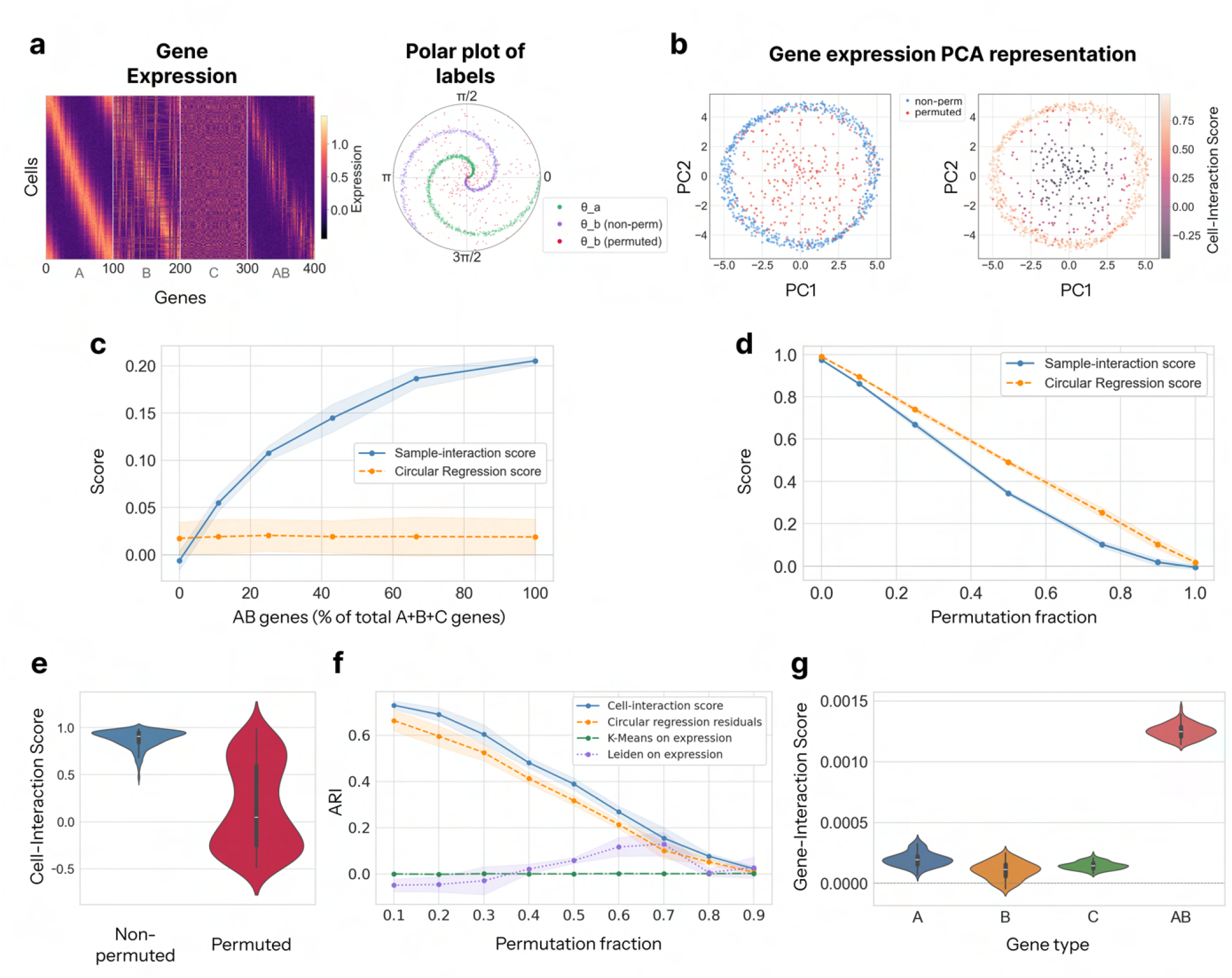
Evaluating CRISP scores on simulated data. **a** Simulated data from two coupled cyclic processes with phases *θ*_*a*_ and *θ*_*b*_ (fixed random offset), with a cellular permutation fraction of 0.3. *Left:* gene expression heatmap of 400 genes (blocks *A, B, C, AB*), cells sorted by *θ*_*a*_. *Right:* each point represents a cell, showing its phases *θ*_*a*_ (green) and *θ*_*b*_ (purple), with permuted cells in crimson. **b** PCA representation of corresponding gene expression. Cells are colored by permutation label (left), and cell-interaction score (right). **c** Sample-interaction score and circular regression score as a function of the number of AB genes. Cell permutation fraction =1. **d** Sample-interaction score and circular regression score as a function of cell permutation fraction. The number of AB genes is fixed at zero. **e** Violin plots of cell-interaction scores for the two cell categories: non permuted (left) and permuted (right). Cell permutation fraction = 0.3. **f** ARI score between the grouping of cells to permuted and non-permuted and clustering based on cell-interaction score, circular regression residuals and two additional partitions of the cells based on K-Means and leiden clustering, as a function of cellular permutation fraction. **g** Violin plots showing gene-interaction score distributions for each gene category (cell permutation fraction = 1).

Next, we assessed whether the CRISP cell-interaction score can distinguish which cells reflect the *A*-*B* coupling. We fixed the number of interacting genes *G*_*AB*_ and varied the permutation fraction *p* ∈ {0.1 … 1}, treating the permuted cells as the ground truth non-interacting population. As a baseline, we used the per-cell residual from the circular regression between *θ*_*a*_ and *θ*_*b*_, which measures the level of deviation of each cell from the global relationship *θ*_*b*_ = *θ*_*a*_ + *ϕ* (Methods 4.6.3). To quantify performance, we clustered cells into two groups by applying K-Means (*k* = 2) to both the per-cell residuals and the CRISP cell-interaction scores, and measured the agreement between each of these partitions and the ground-truth interacting/non-interacting labels using the adjusted Rand index (ARI). Across all permutation levels, the CRISP cell-interaction score consistently outperformed the residual-based baseline (Figure 2f), as well as K-Means and Leiden clustering (*k* = 2) of the expression data.

Finally, we evaluated the CRISP gene-interaction score. When the permutation fraction is set to 1 the two process labels are independent, however, the *AB* genes still co-encode both signals, so the interaction is now driven exclusively by these genes. The *AB* genes were assigned significantly higher CRISP gene-interaction scores than the *A, B*, and *C* groups, enabling localization of the interaction to its driver genes (Figure 2g; Methods 4.6.4).

### 2.3 Dissecting disease-associated zonation interactions in the liver

Liver zonation is the spatial organization of hepatocyte function along the portal-to-central axis of liver lobules, where distinct metabolic and transcriptional programs are prominently expressed in specific zones and the activity of some varies monotonically across the lobule axis [11, 2]. Hepatocytes near the portal vein and those near the central vein are exposed to different oxygen, nutrients, hormones, and morphogens levels, and express distinct sets of genes governing processes such as gluconeogenesis, *β*-oxidation, acid synthesis, and metabolism [11]. We use liver zonation, reconstructed based on the expression of central and portal landmark genes [11], as a test case for CRISP in a pathological context, where zonation–position coupling is disrupted by fibrosis.

#### 2.3.1 Loss of interaction between zonation patterns and physical position in liver fibrosis

In healthy liver, the zonated gene expression of hepatocytes is tightly coupled to their position along the lobule portal-to-central axis [11]. However, in liver fibrosis, marked by excessive extracellular matrix deposition and extensive tissue remodeling [32], as well as vascular distortion, inflammation, and disrupted oxygen and signaling gradients that interfere with positional cues that maintain zone-specific cell identities [2, 10], we expect weakened zonation-position coupling. We used CRISP to characterize the coupling between the zonation signal and physical location of hepatocytes along the progression of fibrosis. To do so, we analyzed a spatial transcriptomics dataset tracking the progression of fibrosis in liver lobules [32], profiling ∼ 300 genes per sample using MERFISH technology (Figure 3a, Methods 4.7.1). We first constructed a cell–cell similarity kernel for each of the two signals, zonation and physical location, using the *k*-nearest-neighbour scheme for continuous signals (Methods 4.3). Next, we quantified the sample zonation–position interaction across the 3 healthy and 4 fibrotic spatial transcriptomics liver samples. Healthy samples received consistently higher zonation-position sample-interaction scores relative to the fibrotic samples (Figure 3b), reflecting a weakening of zonation–position coupling in fibrosis, as expected.

**Figure 3:**
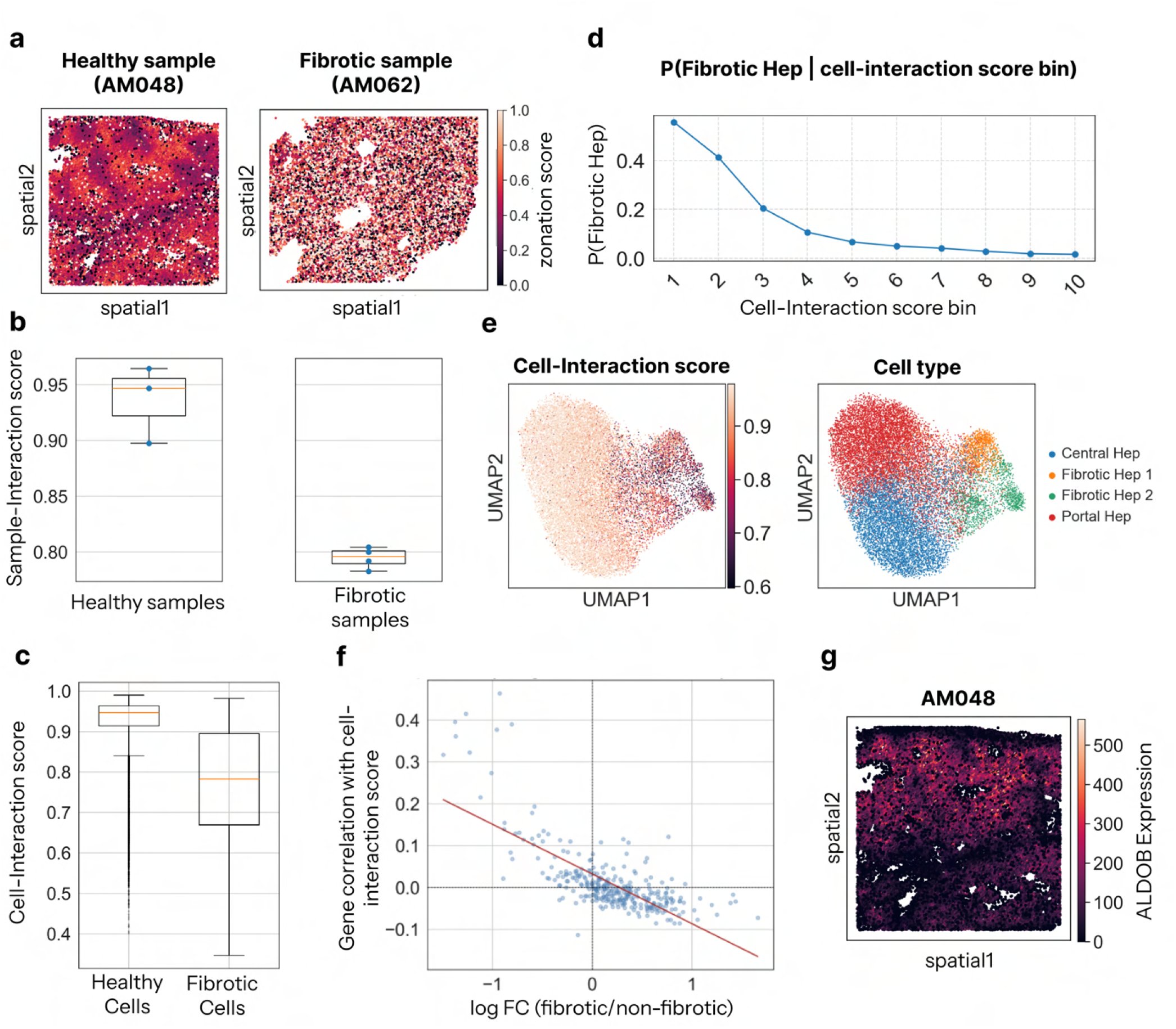
CRISP characterizes fibrotic hepatocytes. **a** Spatial scatter plot visualization of healthy (AM048) and fibrotic (AM062) liver tissues from [32]. Cells are colored according to their zonation score: a continuous portal-central zonation coordinate computed from the summed expression of portal versus central landmark genes (see Methods 4.7.1). **b** Box plots comparing samples-interaction scores for three healthy and four fibrotic samples. **c** Cell-interaction score of healthy versus fibrotic hepatocytes. Middle line in box plot, median; box boundary, IQR; whiskers, 1.5×IQR; minimum and maximum, not indicated in the box plot; black dots, points beyond the minimum or maximum whisker. **d** The probability of a cell to be fibrotic as a function of its cell-interaction score. Score range was divided into ten bins. **e** UMAP representation of the cellular gene expression colored according to their corresponding cell-interaction score (left) and cell type (right). The color scale is clipped at the 2.5th and 97.5th percentiles of the cell-interaction score **f** Each point represents a gene, positioned by its log fold change between fibrotic and non-fibrotic hepatocytes (*x*) and its expression level Pearson correlation with the cell-interaction score across non-fibrotic cells (*y*). Red line is a least-squares linear regression fit, Pearson correlation r = -0.727 (p=2.5e-53). **g** Spatial representation of a healthy tissue. Cells are colored according to their ALDOB expression (highest ranking gene in term of its gene-interaction score). Results in (c-g) are shown for sample AM048. See Supplementary Figure 1 for results on additional samples.

We next used CRISP to localize the disruption in the zonation-position interaction. Each sample, whether annotated as healthy and diseased, comprises both fibrotic and non-fibrotic hepatocytes; hepatocytes were originally annotated by clustering their gene expression into four populations [32], of which we consider “Fibrotic Hep 1” and “Fibrotic Hep 2” as fibrotic, and the zonated “Central” and “Portal” hepatocytes as healthy. Healthy hepatocytes attain higher cell-interaction scores than fibrotic hepatocytes (healthy: median = 0.95, IQR=[0.914, 0.963]); fibrotic: median = 0.78, IQR=[0.669, 0.895]; Figure 3c, e). This contrast extends beyond the binary label: the probability of a cell being fibrotic decreases monotonically with its cell-interaction score (Figure 3d). This trend was consistent across all samples, although it was more pronounced in healthy samples where the zonation–position sample interaction score is consistently higher ([0.89, 0.94, 0.96]) than in fibrotic samples ([0.79, 0.80, 0.79, 0.78]) (see Figure 3 for results on a representative sample, and Supplementary Figure 1 for additional samples).

We next examined whether the cell-interaction score resolves heterogeneity within the healthy hepatocytes population that the binary healthy/fibrotic annotation cannot capture. We correlated the expression of each gene with the cell-interaction score across non-fibrotic cells, and compared this to its differential expression (mean log fold change) between fibrotic and non-fibrotic hepatocytes. We found a significant negative relationship: the downregulated genes in fibrosis tend to be positively correlated with the cell-interaction score in healthy cells (Pearson correlation r = -0.727, p=2.5e-53; Figure 3f). In other words, within the healthy population, a low cell-interaction score corresponds to reduced expression of genes associated with the healthy state, reflecting a potential shift in expression toward the fibrotic state. Together, these findings suggest that the cell-interaction score reflects a continuous gradient of fibrotic transition, with low-scoring healthy hepatocytes representing candidate cells undergoing early or partial fibrotic transition.

Beyond cell-level characterization of the position-zonation interaction, CRISP can be used to identify gene groups which contribute to the interaction i.e. genes that encode both spatial position and zonation. In healthy tissue, where zonation tracks physical position, the expression levels of genes that carry the zonation signal also vary with position, and therefore zonation markers are expected to receive high gene-interaction scores in that case. Indeed, in sample AM048, for example, the genes receiving the highest gene-interaction scores were *ALDOB, ADH1B, FGA, SERPINA1, AHSG, CYP2A6, CYP2E1, EPHX1, ASS1*, and *SLC7A2*, covering both ends of the lobule axis, from pericentral markers like *CYP2E1* to periportal markers like *ASS1, SERPINA1* and *CYP2A6* [32], as well as the periportal-to-mid-lobular markers *ALDOB* and *ADH1B* [32]. Their spatial maps show clear zonal gradients (Figure 3g and Supplementary Figure 2).

### 2.4 Quantifying the Effect of TGF-*β* Dose Driving the Epithelial to Mesenchymal Transition

A perturbation such as TGF-*β* engages several cellular processes at once: it drives cells through the epithelial-to-mesenchymal transition (EMT) while arresting the cell cycle. Therefore we can use CRISP to study which cells are most associated with that interaction. For that, we analyzed a previously collected scRNA-seq dataset of human mammary epithelial (MCF10A) cells undergoing epithelial-to-mesenchymal transition (EMT) [23]. Cells were measured after 14 days of treatment by eight TGF-*β* concentrations (0, 12.5, 25, 50, 100, 200, 400, and 800 pM) (Methods 4.7.2). We first analyzed the interaction between EMT progression and TGF-*β* input, as TGF-*β* is well established as a driver of EMT [21, 23]. EMT progression was represented by the continuous 76GS score [6, 5] using an RBF kernel (see Methods 4.3, 4.7.2), while the EMT signal was represented using a KNN similarity kernel. Analysis of the interaction between TGF-*β* dose and EMT progression with CRISP revealed that the interaction is localized to epithelial cells, with epithelial-like cells receiving substantially higher interaction scores than mesenchymal-like cells (Figure 4f). This result is reflected by the label structure: untreated cells form a tight epithelial group, while almost all treated cells collapse onto the same mesenchymal endpoint (Figure 4e). Because treated mesenchymal cells span many doses, their EMT neighborhoods and dose neighborhoods have a minor overlap, and their cell-interaction scores are low. Furthermore, the low-scoring cells occupy a distinct cell-cycle state, heavily enriched for cells in G1 phase (Figure 4g), matching the known G1 accumulation of terminal mesenchymal cells after prolonged TGF-*β* treatment [20].

**Figure 4:**
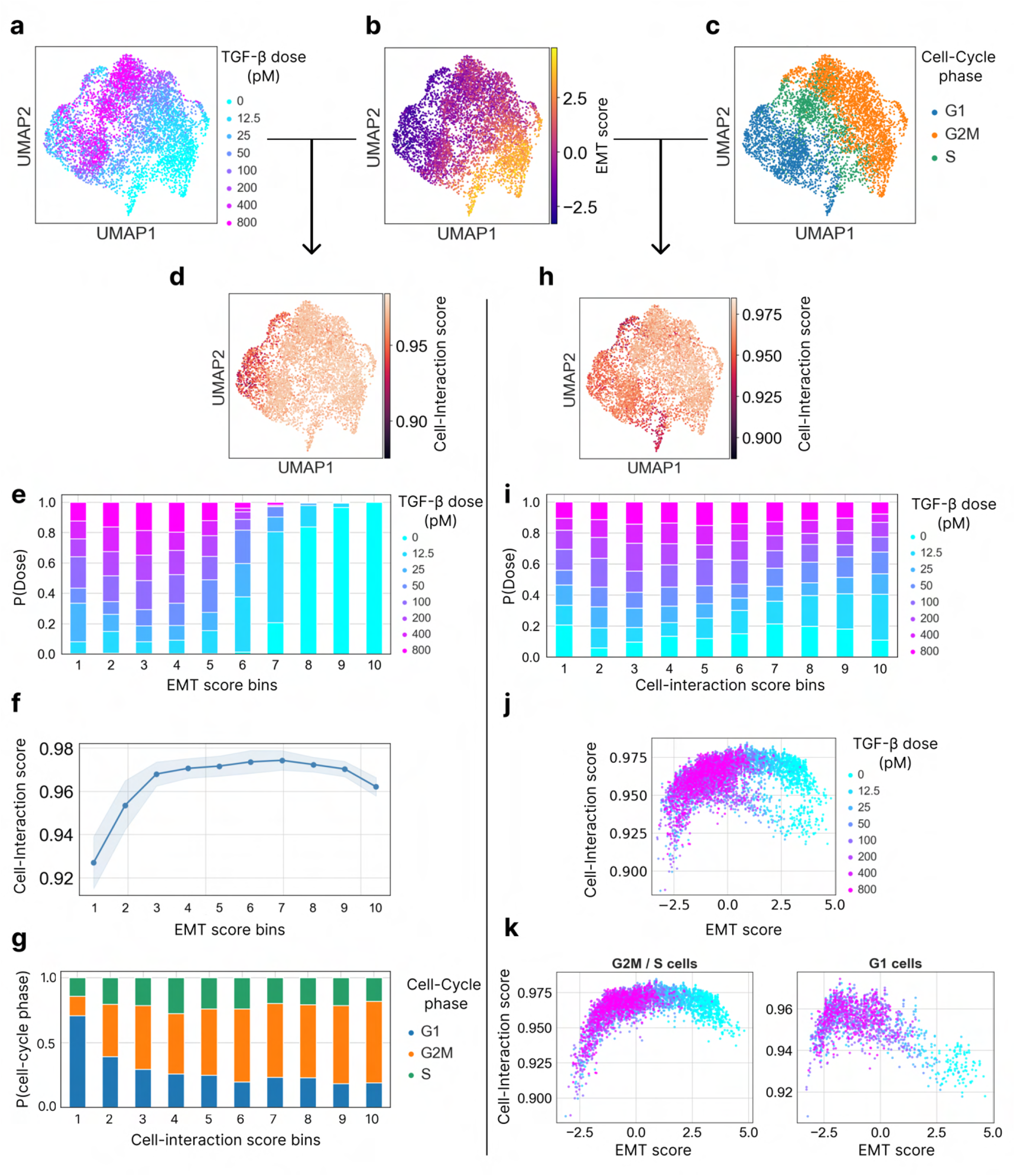
Characterizing interactions between the EMT, TGF-*β* response, and cell cycle progression. **a-d, h** UMAP representation of the TGF-*β*-induced epithelial–mesenchymal transition single-cell dataset from [23], cells colored by (a) TGF-*β* dose, (b) EMT score, (c) cell cycle phase, (d) EMT/TGF-*β* cell-interaction score, and (h) EMT/Cell-Cycle cell-interaction score. **e, i** TGF-*β* dose distribution as a function of (e) EMT score and (i) EMT/Cell-Cycle cell-interaction score; score range was divided into ten equal bins. **f** EMT/TGF-*β* cell-interaction scores as a function of EMT score. Cell-interaction score was binned similarly to (e). **g** Cell-Cycle phase distribution as a function of EMT/TGF-*β* cell-interaction score. **j, k** EMT/Cell-Cycle cell-interaction score as a function of EMT score, colored by TGF-*β* dose, shown for (j) all cells, (k, left) G2M/S cells, and (k, right) G1 cells.

The interaction between EMT and cell-cycle progression (cell-cycle labels for phases G1, S, G2/M, inferred using Scanpy [33] and represented using RBF kernels; Methods 4.3) is lost in a different set of cells than the dose-EMT interaction; Specifically, the EMT-cell-cycle interaction is present in cycling cells and absent in arrested ones. Cells in the S and G2/M phases receive high EMT-cell-cycle cell-interaction scores, while cells in the G1 phase score low and variably (Figure 4h,k). The low-scoring cells originate almost entirely from the highest TGF-*β* doses (Figure 4i,j).

These high-dose cells sit in a stable G1-arrested mesenchymal state where EMT no longer tracks the cell cycle [20]; cells that keep cycling hold onto the EMT–phase coupling at any dose.

### 2.5 Quantifying the interaction between infection time and parasite developmental pseudotime in Plasmodium-infected hepatocytes

Prior to the onset of symptomatic malaria, Plasmodium parasites undergo a developmental phase within hepato-cytes [12, 26]. Development proceeds asynchronously, so cells sampled at the same time following infection span a range of stages. This allowed Afriat et al. [1] to infer a pseudotime trajectory based on the transcriptomes of infected hepatocytes in a single-cell atlas of Plasmodium liver-stage infection, where mouse hepatocytes were sampled across a time course from 2 to 36 hours post-infection (hpi) (Methods 4.7.3). We first aimed to use CRISP to identify cells undergoing delayed parasite development (Figure 5a,b). To this end, we provide CRISP with each cell’s inferred pseudotime and its real infection time (hpi), and encode each using RBF kernels as cell-cell similarity matrices (Methods 4.3).

**Figure 5:**
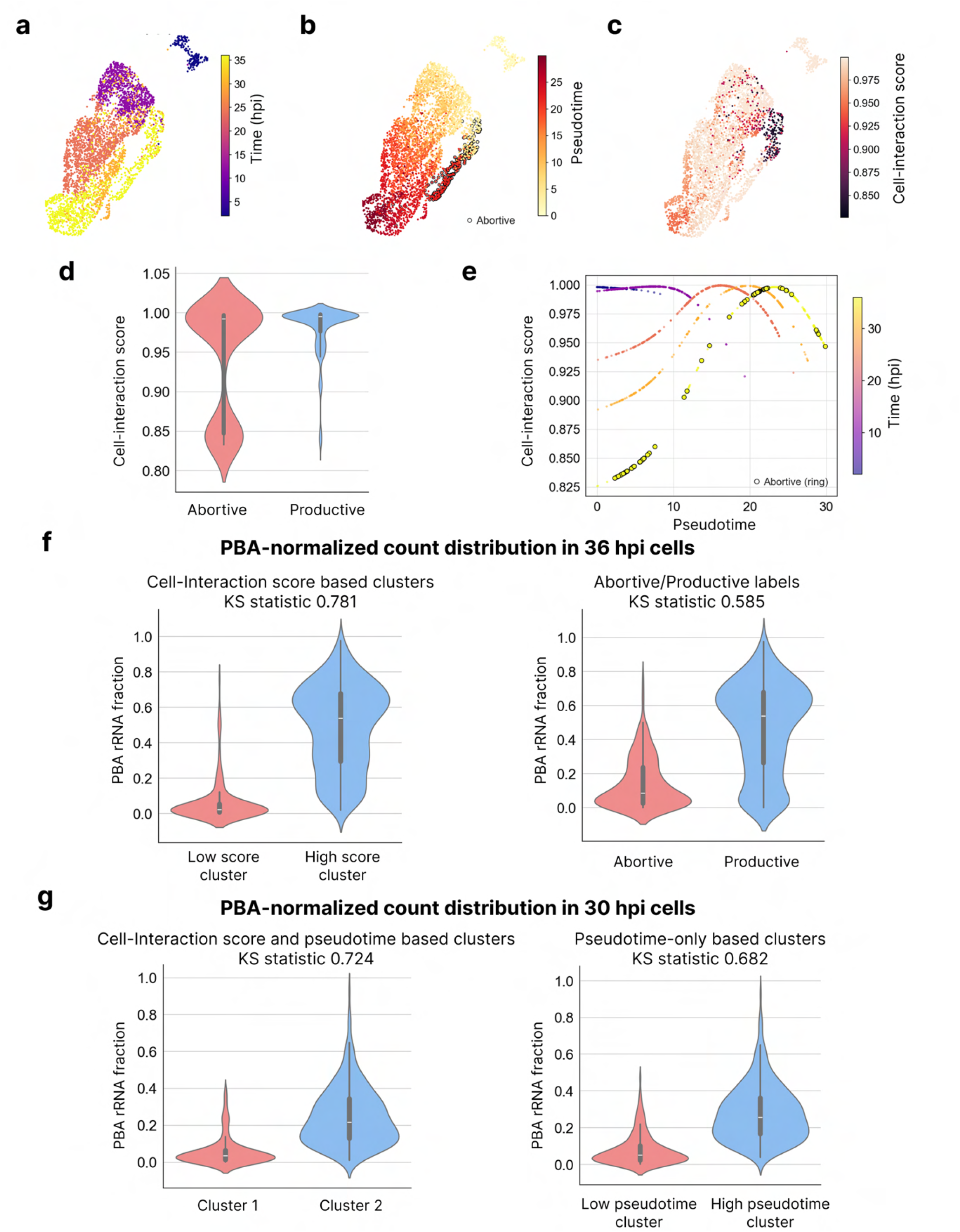
CRISP reveals hepatocytes with delayed parasite development in Plasmodium infections. **a-c** UMAP representation of hepatocytes in a plasmodium-infected liver single-cell dataset [1] colored by real time (a), inferred pseudotime, where abortive-labeled cells are outlined (b), and real time - pseudotime CRISP cell-interaction score (c). **d** Violin plot showing the distribution of CRISP cell-interaction scores in abortive and productive hepatocytes. **e** Scatter plot of pseudotime versus CRISP cell-interaction score, with cells colored by real time; abortive cells are outlined. **f** Violin plots showing normalized parasite burden (PBA) distribution of 36hpi cells clustered according to cell-interaction score (left) and according to productive/abortive labels (right). **g** Violin plots showing normalized parasite burden distribution of 30hpi cells clustered according to cell-interaction score and pseudotime (left) and according to pseudotime only labels (right).

At 36 hpi, Afriat et al. annotated infected hepatocytes as either “productive” or “abortive”, where the latter indicates cells with reduced parasite transcript abundance and signs of failed development [1]. While productive cells scored uniformly high in terms of their pseudotime-real time CRISP cell-interaction scores, as expected, abortive cells exhibited a bimodal distribution, resolving the abortive population into two subpopulations (Figure 5c-e). The low-scoring group of cells undergo delayed parasite development, their low pseudotime is inconsistent with their late (36 hpi) sampling time, and hence these cells receive low cell-interaction scores. In the high-scoring group, pseudotime aligns with the late sampling time, as in productive cells, so the two signals remain coupled and the cells score like productive ones (Figure 5d,e). Thus, low cell-interaction scores appear to be a better marker of cells whose transcriptional progression has become decoupled from real infection time, relative to “productive” / “abortive” annotation scheme.

To test this claim against a molecular measure of infection level, we used the PBA-normalized count, the fraction of Plasmodium transcripts in a cell, as a proxy for parasite developmental state that has been previously used to validate cell-state inference [24, 1]. The CRISP cell-interaction score measures how well a cell’s pseudotime aligns with its sampling time; it is not, in general, a measure of parasite load. At 36 hpi, however, when the cells are parasite-rich, the two coincide, and a low score then marks a lagging cell with low parasite load. Indeed, clustering cells by their cell-interaction score provided a better separation of the PBA distributions compared to the 36 hpi abortive/productive annotation (Kolmogorov–Smirnov statistic 0.781 versus 0.585; Figure 5f), demonstrating that identification of cells with delayed parasite development could be done more faithfully by the cell-interaction score than the original labels.

At the intermediate 30 hpi time point, which carries no abortive/productive annotation, previous smFISH analyses [1] nonetheless suggested that some cells are already developmentally delayed and we next studied whether the cell-interaction score could be used to identify them. Clustering cells by their assigned pseudotime separates the PBA distributions, and hence distinguishes between different developmental states (KS statistic 0.682; Figure 5g – right). Adding the CRISP cell-interaction score as a second clustering feature sharpened this separation (KS statistic 0.724; Figure 5g – left), improving the estimate of parasite developmental state beyond pseudotime alone. Together, these results suggest that the CRISP cell-interaction score captures biologically meaningful heterogeneity beyond the existing binary labeling scheme.

## 3 Discussion

We introduced CRISP, a framework for quantifying and localizing interactions between different spatial-temporal signals encoded in single-cell and spatial omics data. CRISP characterizes signal-interactions at the sample-level and guides the localization of signal interactions, identifying the cellular subpopulations that drive or disrupt global coupling between biological signals, and pinpoints the genes that jointly encode both signals, offering a candidate list of shared regulators. Because CRISP reads interaction from expression rather than labels, it detects coupling that labels-based tests miss: two signals can share a gene-expression encoding even when their labels are statistically independent. We validated CRISP on simulations of two interacting cyclic processes with tunable coupling, and demonstrated its utility on three single-cell and spatial omics datasets. In liver fibrosis, CRISP recovered the loss of zonation–position coupling at the sample, cell, and gene levels. In TGF-*β* induced EMT, CRISP resolved the asymmetric coupling between dose, EMT progression, and cell-cycle phase. In Plasmodium liver-stage infection, CRISP localized transcriptionally delayed abortive hepatocytes and characterized this cellular state more faithfully than existing labeling schemes.

CRISP quantifies a specific operational notion of interactions: two signals are modeled as interacting when the expression patterns they induce are correlated following projection onto their respective kernels. Other forms of signal interactions may be explored in future work. Additionally, the interactions detected by CRISP are statistical rather than causal, and indicate that two signals share a gene-expression encoding, but cannot distinguish, for example, direct coupling from co-regulation by a common upstream factor, nor establish directionality. A further limitation is that errors in the input labels, such as inferred pseudotime or cell-cycle phase, may propagate into the interaction scores.

We see several directions for future work. Extending the CRISP framework to quantify higher-order interactions could reveal coupling between multiple multicellular signals, such as joint encoding of space, time, and perturbation. In addition, we envision that CRISP could be embedded within a generative model, enabling counterfactual analysis of how intervening in one signal reshapes the expression associated with the others.

## 4 Methods

### 4.1 The CRISP workflow

Given input data consisting of a single-cell or spatial omics gene expression matrix *X* ∈ ℝ^*N* ×*G*^ (*N* cells, *G* genes) and two cell-level signals 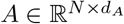 and 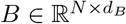, where *d*_*A*_ and *d*_*B*_ are the dimensions of the respective label spaces, the CRISP workflow is as follows:

1. Preprocess the data (see Methods 4.7).
2. Generate signal-specific cell–cell similarity kernel for each signal *K*_*A*_ ∈ ℝ^*N* ×*N*^ and *K*_*B*_ ∈ ℝ^*N* ×*N*^ , representing its underlying topology (see Methods 4.3).
3. Project the expression matrix onto each kernel to obtain the signal-associated components *X*_*A*_ = *K*_*A*_*X* and *X*_*B*_ = *K*_*B*_*X*.
4. Compute the cell-interaction score CRISP_cell_(*i*), for each cell *i*, as the Pearson correlation between the corresponding rows *X*_*A*_[*i*, :] and *X*_*B*_[*i*, :], taken across genes.
5. Average the cell-interaction score to get a single sample-interaction score CRISP(*A, B*).
6. For each gene *g*, compute its gene-interaction score CRISP_gene_(*g*) as the contribution of gene *g* to the cell-interaction score, averaged over all cells (Methods 4.5).

### 4.2 Rationale

The aim of CRISP is to quantify the extent to which two signals are jointly encoded in a multicellular population, measured via a gene expression matrix. Our underlying assumption is that two signals interact when the gene-expression patterns they induce share a common structure, that is, when the component of expression attributable to one signal is similar to the component attributable to the other.

CRISP therefore proceeds in two steps: it first isolates the component of gene expression that varies with each of the two signals, and then measures how similar these two components are. The first isolation step is performed as a probabilistic filtering based on the SiFT algorithm [25]: a signal-specific cell-cell similarity kernel *K* assigns high similarity to pairs of cells that are similar with respect to that signal alone. The projection *KX* replaces each cell’s expression by the similarity-weighted average over its neighbors, retaining the part of expression that co-varies with the signal. Whereas SiFT subtracts *KX* from *X* to remove a signal, CRISP keeps *KX* as the signal-specific component.

Applying this to both signals yields *X*_*A*_ = *K*_*A*_*X* and *X*_*B*_ = *K*_*B*_*X*. If signals *A* and *B* are encoded by overlapping gene expression programs, *X*_*A*_ and *X*_*B*_ will be correlated.Computing the correlation between the rows of *X*_*A*_ and *X*_*B*_ results in the cell-level score from which both the global and gene scores are derived 4.5.

### 4.3 From signals to kernels

To extract the component of the data associated with each signal, we construct a signal-specific cell–cell similarity kernel that represents the signal, where they are first mapped to a Euclidean embedding space (4.7). Each embedding is converted into a row-stochastic kernel *K* ∈ [0, 1]^*N*×*N*^ , whose row *i* defines a probability distribution *p*_*i*_(·) over cells, assigning higher weight to cells close to cell *i*. CRISP handles signals defined both on a discrete or continuous domain.

#### Continuous signals

When the signal is defined on a continuous domain (e.g. spatial position), CRISP constructs a *k*-nearest-neighbour graph on the signal’s embedding (*k* = 15) using the standard ScanPy construction (sc.pp.neighbors) [33]: for each cell *i*, the *k* nearest neighbours are assigned connectivity weights that decrease with distance, where all other cells are assigned a zero weight. Each row is then normalized by its sum,

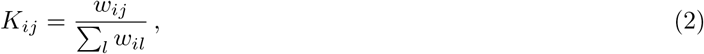

where *w*_*ij*_ is the connectivity weight between cells *i* and *j*, so that row *i* defines a probability distribution *p*_*i*_(·) over cells, concentrated on cell *i*’s neighbourhood.

#### Discrete signals

When the signal is defined on a discrete domain (e.g. cell type), many cells share identical signal values. Therefore, we replace the *k*-nearest-neighbour graph with a more informative representation of data similarity structure for that setting via an RBF kernel:

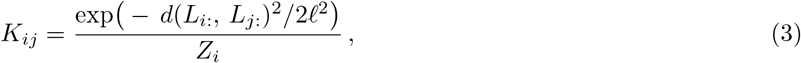

where *l* is a length-scale controlling the effective radius of cells that have a non-negligible influence, set by default to half the median pairwise d istance; *d*(*·, ·*) is the Euclidean distance; *L*_*i*:_ is the label (or its encoding, see Methods 4.7) of cell *i*; and *Z*_*i*_ = ∑_*j*_ exp (− *d*(*L*_*i*:_, *L*_*j*:_)^2^*/*2*l*^2^) is a per-row normalization factor.

### 4.4 Projected expression and its stochastic interpretation

For a kernel *K*, each entry of the projected expression matrix *P* = *KX* can be written as

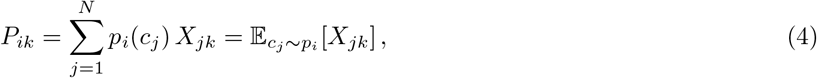

where *p*_*i*_(*c*_*j*_) denotes the probability distribution for cell *i* and *X*_*jk*_ the expression level of gene *k* in cell *j*.

Therefore, the projected expression of gene *k* in cell *i* is the expected expression under the similarity-weighted neighborhood distribution of cell *i. P* is thus a smoothed version of *X* in which each cell is represented by the expectation of expression over its neighborhood. This projection step is inspired by the SiFT algorithm [25], which subtracts *KX* from *X* to remove, or filter, a target signal; CRISP instead retains *KX* as the signal-specific component.

For the two input signals, CRISP computes

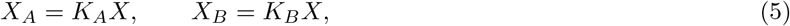

interpreted as the components of expression associated with signals *A* and *B*, respectively.

### 4.5 Interaction scores

- **Cell-interaction score:** for cell *i*, CRISP_cell_(*i*) is the Pearson correlation between rows *X*_*A*_[*i*, :] and *X*_*B*_[*i*, :] across the *G* genes:

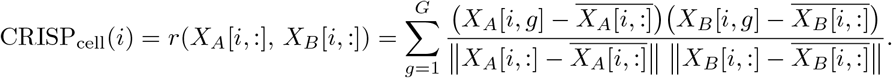
- **Sample-interaction score:** The mean of the cell-interaction score yielding a dataset-wide interaction value between signals *A* and *B*:

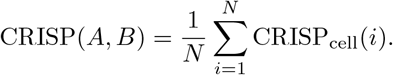
- **Gene-interaction score:** the cell-interaction score decomposes additively over genes, so gene *g*’s contribution to cell *i*’s score is

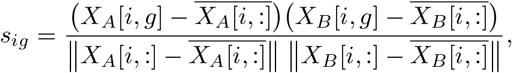

and the gene-interaction score is the mean of this contribution over cells, 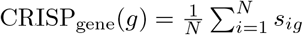.

### 4.6 Simulation

#### 4.6.1 Simulated data

Each synthetic dataset contains *N* = 1000 ordered cells. Three cyclic labels are assigned per cell: *θ*_*a*_ value at cell i is given by *θ*_*a,i*_ = 2*πi/n, θ*_*b*_ = (*θ*_*a*_ + *φ*) mod 2*π*, where *φ* ∼ Unif[0, 2*π*) is a single random phase fixed per dataset, so processes *A* and *B* are initially fully coupled (their phase difference *θ*_*b*_ −*θ*_*a*_ ≡ *φ* is constant across all cells), and *θ*_*c*_ is obtained by randomly shuffling the same phases {2*πi/N*}_*i*_ across cells, independently of *θ*_*a*_ and *θ*_*b*_. The expression of each gene is then generated using a periodic kernel applied to its label value (*θ*_*a*_, *θ*_*b*_ or *θ*_*c*_): for example, each A-gene *g* is assigned a peak *µ*_*g*_ ∼ Unif[0, 2*π*) and its expression level in cell i is given by 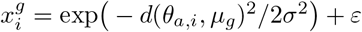, with *d*(*α, β*) = min(|*α* − *β*|, 2*π* − |*α* − *β*|), *σ* = *π/*4 and *ε* ∼ *N* (0, 0.1) an additive Gaussian noise. B- and C-genes are simulated similarly based on *θ*_*b*_ and *θ*_*c*_, respectively.

To weaken the coupling, we can vary the first parameter, the cell permutation fraction *ρ*. A subset *P* of ⌊*ρN*⌉ cells is sampled uniformly at random, and their *B*-gene rows and *θ*_*b*_ values are jointly shuffled by a random permutation

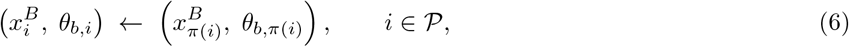

where 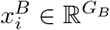 corresponds to the expression levels of B-genes in cell *i*, and *θ*_*b,i*_ is the *θ*_*b*_ value of cell *i*. Cells in the subset *P* are labeled as permuted.

The second parameter is the number of interacting genes, *G*_*AB*_; following the cell permutation step, each AB gene g is simulated as the product of two periodic kernels, such that g expression level in cell i is given by 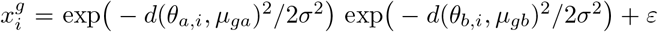 with *µ*_*ga*_, *µ*_*gb*_ ∼ *U* (0, 2*π*) independently, *σ* = *π/*4 and *ε* ∼ *N* (0, 0.1) an additive Gaussian noise. Generated this way, *g* hence encodes both signals. These AB genes mirror the multiplicative *Z* × *R* genes modeled in [8]. The final expression profile of each cell is the concatenation of the four blocks

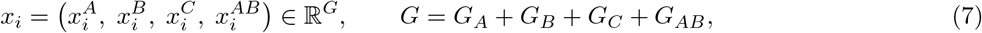

##### Signal Encoding

In all simulation experiments, we use cyclic encoding of angles: for each pair of cells (*u, v*) we compute the circular distance *d*_*uv*_ = min(|*θ*_*u*_ − *θ*_*v*_|, 2*π* − |*θ*_*u*_ − *θ*_*v*_|) and define the affinity matrix 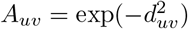.

The matrix encodes the circular topology of the data and is passed as input to the cell-cell similarity kernels.

#### 4.6.2 Sample-interaction score experiment

We fixed a pool of *G*_*A*_ = 100 *A*-genes, *G*_*B*_ = 100 *B*-genes, and *G*_*C*_ = 100 background genes. In the first scenario, we checked the influence of AB genes on both the circular-regression baseline and the CRISP sample-interaction score: we varied the relative ratio of interacting genes *G*_*AB*_*/*(*G*_*A*_ + *G*_*B*_ + *G*_*C*_) ∈ {0, 20, 40, 60, 80, 100} with permutation fraction p=1 and 5 random seeds per cell. In the second scenario, we assessed the influence of the permutation fraction on both methods: we fixed the relative ratio of interacting genes at 0 and varied the permutation fraction *ρ* ∈ {0, 0.1, … , 0.9}. For each dataset we recorded the sample-interaction score and a circular-regression baseline measuring how closely the labels follow 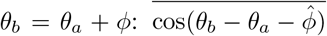 with 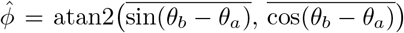 [19]. Scores were then averaged over seeds.

#### 4.6.3 Cell-interaction score experiment

To evaluate the cell-interaction score we fixed the number of interacting genes at *G*_*AB*_ = 0 and swept the permutation fraction *p* ∈ {0.1, … , 1} on 100 *A*-genes, 100 *B*-genes, and 100 *C*-genes, with 5 seeds. We considered non-permuted cells as interacting, and permuted cells as non-interacting. For both the cell-interaction score and the baseline circular-regression residual 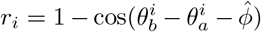, we clustered the per-cell scores into two groups using k-means (k=2) and quantified overlap with the ground-truth permuted/non-permuted partition with adjusted Rand index (ARI); across all permutation levels the cell-interaction score outperformed the residual baseline. As a control, we also partitioned the cells with both *k*-means and Leiden clustering on the full expression matrix; performing the same comparison with ground-truth, both yielded ARI near zero, confirming that the interacting/non-interacting split is not the dominant structure in the gene expression matrix.

#### 4.6.4 Gene-interaction Score Experiment

We generated a synthetic dataset of 1000 cells with four equal gene blocks (100 A, 100 B, 100 background C, and 100 AB interaction genes; noise *η* = 0.1) with cell permutation fraction *ρ* of 1, so that *A*-*B* interaction originates solely from AB genes. We then computed and visualized the distribution of the gene-interaction score within each gene program as violin plots 2g.

### 4.7 Datasets

All datasets underwent the same standard preprocessing using scanpy [33]: library-size normalization to 10^4^ counts per cell (`sc.pp.normalize total`) followed by log(1 + *x*) transformation (`sc.pp.log1p`), after which dataset-specific steps were applied, as detailed below.

#### 4.7.1 Liver fibrosis

##### Data Preprocessing

In section 2.3.1 we analyzed a MERFISH spatial-transcriptomics dataset of human liver spanning healthy and fibrotic (diseased) donors [32], comprising seven samples (AM031, AM042, AM048, AM061, AM062, AM066, AM072). Each cell carries its spatial coordinates and a curated cell-type annotation. Because the interaction of interest is between hepatic zonation and tissue architecture, we restricted the analysis to hepatocytes, retaining the central, portal, and fibrotic hepatocyte subtypes and discarding all other cell types.

##### Zonation Inference

For each cell we reconstructed a continuous zonation coordinate *µ* along the portal-central axis, zonation_position ∈ [0, 1]. Using curated central and portal landmark genes present in the MERFISH panel, each landmark’s expression was normalized to its maximum across cells and summed within its class; the coordinate was then taken as *µ* = *S*_portal_*/*(*S*_portal_ + *S*_central_), where *S*_portal_ and *S*_central_ are the summed portal and central landmark expressions. This zonation coordinate serves as one of the two interacting signals in the CRISP analysis, along with the spatial position of the cell.

The zonation coordinate was used directly as a 1D continuous signal, and the spatial signal was taken as the measured physical (x,y) position.

#### 4.7.2 TGF-*β*/EMT data

##### Data Preprocessing

In section 2.4 we analyzed a droplet-based scRNA-seq dataset of epithelial cells driven through the epithelial–mesenchymal transition by TGF-*β* stimulation [23]. Starting from the raw count matrix, we kept the top 12,000 most dispersed genes (`sc.pp.filter genes dispersion`), and removed duplicate gene symbols.

##### EMT score inference

First, a continuous EMT score was computed based on the 76-gene signature (76GS) method [5, 6]. Let *G* denote the set of signature genes detected in the dataset and *x*_*ig*_ the expression of gene *g* in cell *i*. Each gene is assigned a weight coefficient *w*_*g*_ = *ρ*(*x*_*g*_, *x*_CDH1_), the Pearson correlation across cells between the expression level of gene *g, x*_*g*_, and that of the epithelial marker *CDH1* (E-cadherin), *x*_CDH1_. The EMT score of cell *i* is computed as the correlation-weighted sum of its signature genes expression, centered across cells:

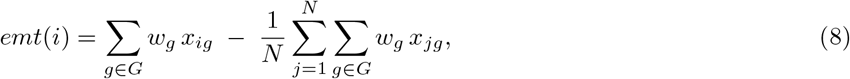

where *N* is the number of cells. Thus, epithelial (mesenchymal) genes, whose expression levels are positively (negatively) correlated with *CDH1* expression, thus increase (decrease) the EMT score and indicate a more epithelial (mesenchymal) state.

##### Signal Encoding

As explained in 4.3, for some non-euclidian topologies, labels may be embedded before kernel computation so that pairwise similarities reflect the underlying label geometry. Here we embed the cyclic cell-cycle phases into a circular affinity matrix as was done for the simulated data (4.6.1).

Since the cell-cycle label has only three values, we set the RBF length scale (Methods 4.3) to a small enough value (*l* = 10^−4^), so that effectively, affinity is nonzero only between cells sharing the same phase. The resulting kernel is block-diagonal, grouping cells by phase.

#### 4.7.3 Plasmodium-infected liver data

##### Data Preprocessing

In section 2.5 we analyzed the *Plasmodium* liver-stage spatiotemporal single-cell atlas of [1] and followed preprocessing steps similarly to [24]: we retained only infected cells and selected the top 2,000 highly variable genes from the raw counts using `seurat v3` [27] (`sc.pp.highly variable genes(flavor=seurat v3`)). The dataset includes, for each cell, both an experimental time given by its hours post-infection (hpi), and an inferred pseudotime capturing parasite progression along the infection trajectory. Both labels were used to construct the corresponding cell-cell similarity kernels. Clustering in downstream analysis was done using K-Means with K=2.

### 4.8 Data availability

All datasets analysed in this study are publicly available. The healthy and fibrotic liver MERFISH dataset is avail-able on the Dryad Digital Repository: https://doi.org/10.5061/dryad.37pvmcvsg. The single-cell RNA-seq dataset of TGF-*β*-induced epithelial–mesenchymal transition was obtained from GEO under accession code GSE213753. The spatio-temporally resolved single-cell atlas of the Plasmodium liver stage is deposited at: https://doi.org/10.5281/zenodo.7081863.

### 4.9 Code availability

All code for Crisp is publicly available on GitHub at https://github.com/nitzanlab/crisp.

## 5 Acknowledgments

We would like to express our gratitude to Guy Pelc for insightful discussions and all members of the Nitzan lab for general feedback. This work was supported by the Abisch-Frenkel Foundation, the Israel Science Foundation (grant no. 1079/21, 1901/26), the European Union (ERC, DecodeSC, 101040660) (M.N.), and by the Israeli Council for Higher Education Ph.D. fellowship (J.K.). Views and opinions expressed are, however, those of the author(s) only and do not necessarily reflect those of the European Union or the European Research Council.

## Supplementary Figures

**Supplementary Figure 1:**
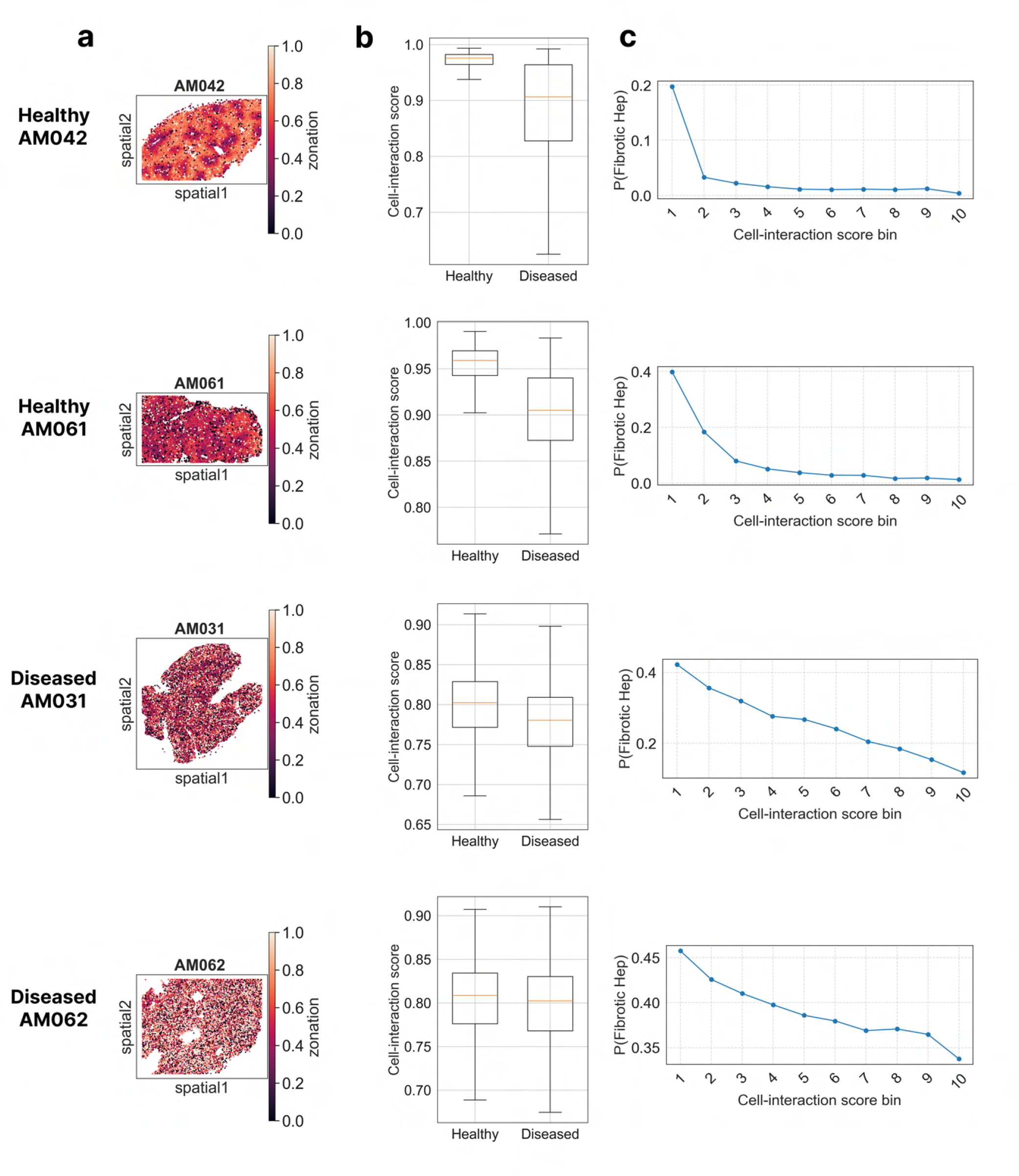
CRISP cell-interaction scores results for liver tissue samples. **a** Spatial scatter plot visualization of two healthy (AM042, AM061) and fibrotic (AM031, AM062) liver tissues from [32]. Cells are colored according to their zonation score: a continuous portal-central zonation coordinate computed based on the summed expression of portal versus central landmark genes (see Methods 4.7.1). **b** Cell-interaction score of healthy versus fibrotic hepatocytes. Middle line in box plot, median; box boundary, IQR; whiskers, 1.5×IQR; minimum and maximum, not indicated in the box plot; black dots, points beyond the minimum or maximum whisker. **c** Plot representing probability of a cells to be fibrotic as a function of its interaction score. Score range was divided into ten bins.

**Supplementary Figure 2:**
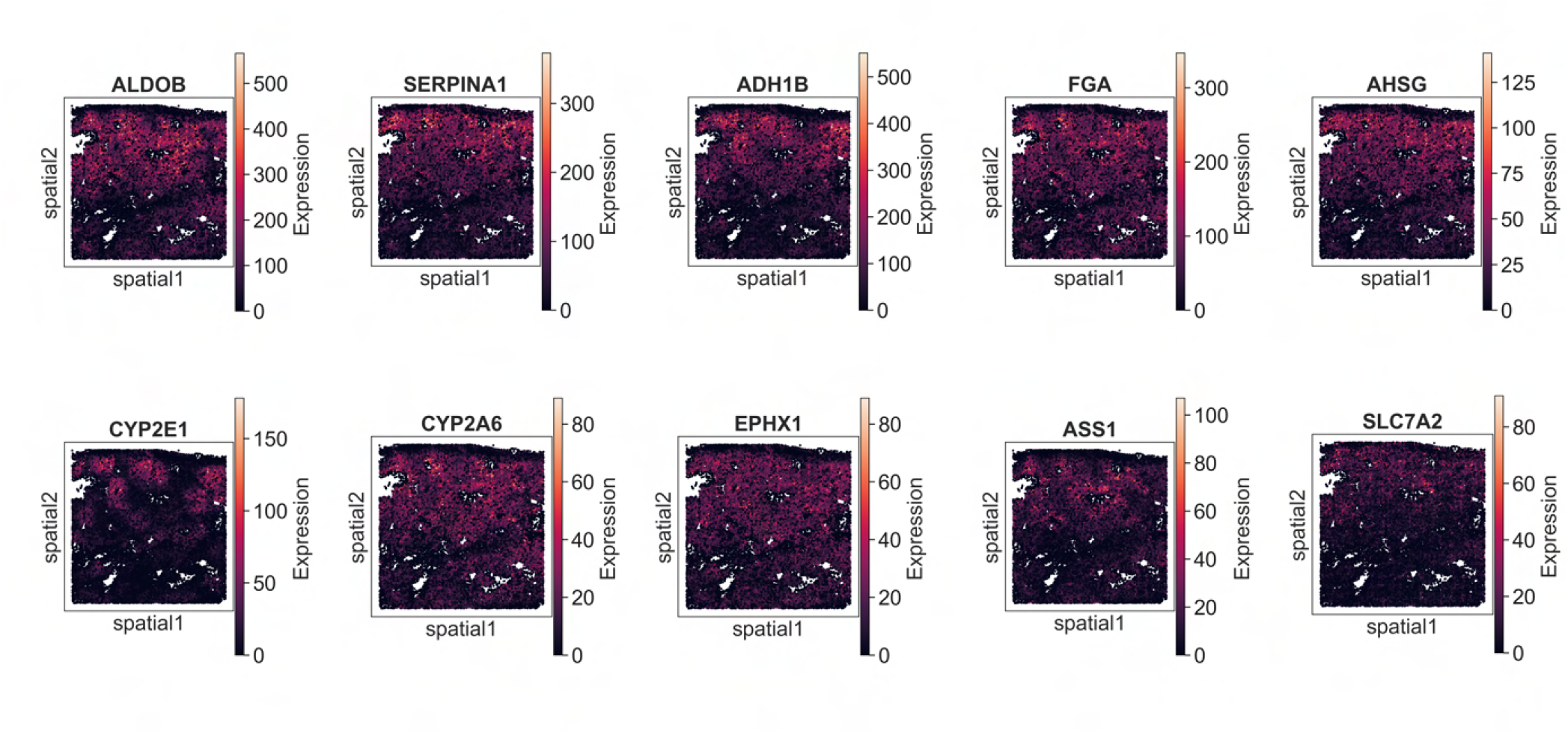
Gene-interaction score reveals zonation markers. Results are shown for a healthy liver tissue sample, AM048, from the spatial transcritomics dataset studied in 2.3.1 from [32]. Cells are colored based on the expression levels of the top 10 genes ranked according to their gene-interaction scores.

